# Exploring the mechanisms and potential therapeutic targets of Ferroptosis Related Genes in ankylosing spondylitis

**DOI:** 10.1101/2023.09.19.558184

**Authors:** Zhen-Gang Liu, Fan Yang, Bing Chen, Peng-Fu Li, Bo-Yin Zhang

**Author notes:** Corresponding Author: Bo-Yin Zhang.

## Abstract

**Background:** Ferroptosis is a novel type of regulated cell death, and there is growing evidence that it is directly associated with the disease. Therefore, this study aimed to investigate the relevance of iron Ferroptosis-related genes to ankylosing spondylitis (AS) to propose a novel targeted therapy for AS patients.

**Methods:** AS samples were downloaded from the Gene Expression Omnibus (GEO) database (GSE41038). Ferroptosis-related genes were obtained from the FerrDb database, and differential expression analysis was performed using the GEO2R tool. Additionally, we constructed a protein-protein interaction (PPI) network and identified the central genes using Cytoscape. A neural network model was developed utilizing the data mining software Clementine. cMap technology was used to screen potential pharmacological compounds that target the central gene,and a predictive model was built by machine learning methods.

**Results:** We discovered 786 differentially expressed genes in AS samples compared to normal tissue, including 359 up-regulated and 427 down-regulated genes. The intersection of ferroptosis-related genes and differential genes yielded 20 overlapping genes. We constructed a hub gene co-expression network with 20 nodes and 14 edges and obtained the ten most recommended drugs for AS.

**Conclusion:** Bioinformatics analysis identified 20 potential genes associated with Ferroptosis in AS. Genes such as CAV1, NOX4, and NQO1 were revealed to influence the development of AS by regulating Ferroptosis. Our study provides new insights into the function of iron Ferroptosis-related genes in AS, suggesting that targeting Ferroptosis may be a potential therapeutic option for AS.

## Introduction

Cell death is necessary for normal development, homeostasis, and the prevention of over-proliferative diseases, including cancer ^1^. Ferroptosis is a novel method of regulating cell death that is highly dependent on lipid peroxidation of iron and is morphologically, biochemically, and genetically distinct from apoptosis, necrosis and autophagy. Ferroptosis involves a variety of biological processes, including iron metabolism, lipid metabolism, oxidative stress, and the biosynthesis of nicotinamide adenine dinucleotide phosphate (NADPH), glutathione (GSH), and coenzyme Q10 (CoQ10) ^2^. Dysregulation of iron metabolism, including redox reactions of iron in the body, can result in excessive production of reactive oxygen species (ROS), and excessive production of ROS can cause damage to genetic material such as DNA and RNA, which is associated with cancer and neurodegenerative diseases. In RAS-mutated cancer cells, transferrin levels are elevated, which increases iron levels within the cells. High iron levels increase the sensitivity of cells to the Ferroptosis inducers erastin and RSL3 ^3^. Iron death plays an important role in a variety of tumor diseases, neurological disorders ^4^, ischemia-reperfusion injury ^5^, liver, kidney ^6^, and other organ-related diseases. In clinical studies, this discovery has contributed to the development of a new cytoprotective strategy, and while initial progress has been made on the mechanism of Ferroptosis, the specific mechanism of Ferroptosis still requires further investigation.

Ankylosing spondylitis (AS) is a chronic immune-mediated inflammatory arthritis, predominantly in men ^7^. Back pain and stiffness typically appear for the first time between 20 and 30 ^8^. It is characterized by inflammation of the mesial skeleton, peripheral joints, and attachment points, with a high genetic susceptibility, including reactive arthritis, enteropathic arthritis, and psoriatic arthritis ^8^. Comorbidities of AS include diseases or manifestations of the skin, eyes, bones, gastrointestinal and genitourinary tracts, heart, lungs, kidneys, and nervous system ^9^. The etiology and pathogenesis of AS remain unknown. It is believed that genetic, infectious, environmental, and immune factors are involved in the pathogenesis of AS. The genetic role in the pathogenesis of AS is the subject of the most intensive research ^10^. Human leukocyte antigen (HLA)-B27 is strongly associated with disease ^11^. Non-steroidal anti-inflammatory drugs (NSAIDs) are the first-line therapy for relieving inflammation. Second-line treatments have had limited success, including corticosteroids and various disease-modifying antirheumatic drugs. In addition to alleviating symptoms, emerging biological therapies may positively modify the disease process.

The role of iron death-associated genes in developing ankylosing spondylitis is poorly understood, and targeting these genes for treating AS has received limited attention. In this study, we obtained clinical data from a public database of ankylosing spondylitis patients. We performed a bioinformatics-based search for iron death-related genes and pathways to identify potential therapeutic agents for AS.

## Materials and methods

### Data collection and preprocessing

In this study, we downloaded a microarray dataset (GSE41038) from the Gene Expression Omnibus (GEO) database (https://www.ncbi.nlm.nih.gov/geo/) to investigate changes in gene expression profiles in patients with Ankylosing spondylitis. The gene dataset GSE41038 ^12^ includes two samples of ankylosing spondylitis and four samples of normal tissue. The Ferroptosis related databaseFerrDb (http://www.zhounan.org/ferrdb/index.html) and the GeneCards database (https://www.genecards.org/) were used to find microarray data and ferroptosis-related genes ^13^. Two ankylosing spondylitis (AS) and four normal control biopsies were obtained from knee joint synovial biopsy samples.

### Identification of DEGs related to ferroptosis

A volcano map was used to describe differential genes using the R software package limma for differential expression analysis with p < 0.05 and |log2 FC| > 1 as filtering conditions. In the map, red nodes represent up-regulated DEGs, blue nodes represent down-regulated DEGs, and gray nodes are genes without differences.

### Ferroptosis-related genes and Venn analysis

We extracted a dataset of 265 genes from the Ferroptosis database and intersected it with GSE41038 to identify ferroptosis DEGs. We overlapped these genes with genes from the ankylosing spondylitis-associated module. Using the online tool Venny2.1 (http://bioinformatics.psb.ugent.be/webtools/Venn/), Venn diagrams of the DEGs were constructed to describe the details of the overlapping genes.

### Gene Ontology Enrichment Analysis and Kyoto Encyclopedia of Genes and Genomes Pathway Analysis

FunRich-3.1.3 tool (http://www.funrich.org/) was used for gene enrichment analysis. Gene ontology (GO) functional enrichment analysis was used for molecular function [MF], biological process [BP], and cellular component [CC]. Kyoto Encyclopedia of Genes and Genome (KEGG) pathway analysis of the DEGs was performed in the Database for Annotation. A P-value < 0.05 was considered statistically significant.

### Protein-Protein Interaction Network Construction and identification of hub genes

The ferroptosis-related PPI network was constructed with STRING (DEGs were analyzed to predict the interaction relationships between proteins encoded by genes that may play a significant role in the pathogenesis of ankylosing spondylitis. The algorithm used to screen the top ten hub genes with CytoHubba ^14^ in Cytoscape was the degree of connectivity.

### Machine learning model building and evaluation

We used machine learning methods to construct prediction models based on the data mining software Clementine and divided the dataset GSE41038 into training and validation sets in a ratio of 3:1. The neural network was constructed with the presence or absence of AS as the output variable and genes as the input variables.

### Screening for Potential Pharmacological Targets

The cMAP (ConnectivityMap) database (https://clue.io/query) includes data on gene expression profiles induced by 33,609 small molecule compounds acting on multiple cell lines. It cand to compare drug-induced gene profiles and gene expression. Screening results of potential pharmacological targets were downloaded from cMAP, ranked and filtered based on drug connectivity scores. Top 10 medications suggested for AS were NCF2, TBX2, PP-30, ADAM15, SLC2A6, HSP90AB1, entinostat, KU-0063794, COB, and ochratoxin-a.

## RESULT

### Identification of Differentially Expressed Genes

The gene dataset GSE41038 consisted of two samples of ankylosing spondylitis and four samples of normal tissues. Using the limma package in the R software package for differential expression analysis and p < 0.05 and |log2 FC| > 1 as filtering conditions, we identified 786 genes differentially expressed in ankylosing spondylitis samples to normal tissues. Among 786 identified genes, 359 were up-regulated, while 427 were down-regulated. Cluster analysis of these genes for differential genes is displayed in the volcano plot (Figure 1A). Data normalization and cross-comparability were also evaluated. The box line plot (Figure 1B) indicates the central distribution of the selected samples, and the numerical distribution follows the criteria, indicating that the microarray data have high quality and cross-comparability. The heatmap of the dataset (Figure 1C) depicts improved sample clustering and higher confidence levels.

**Figure 1.**
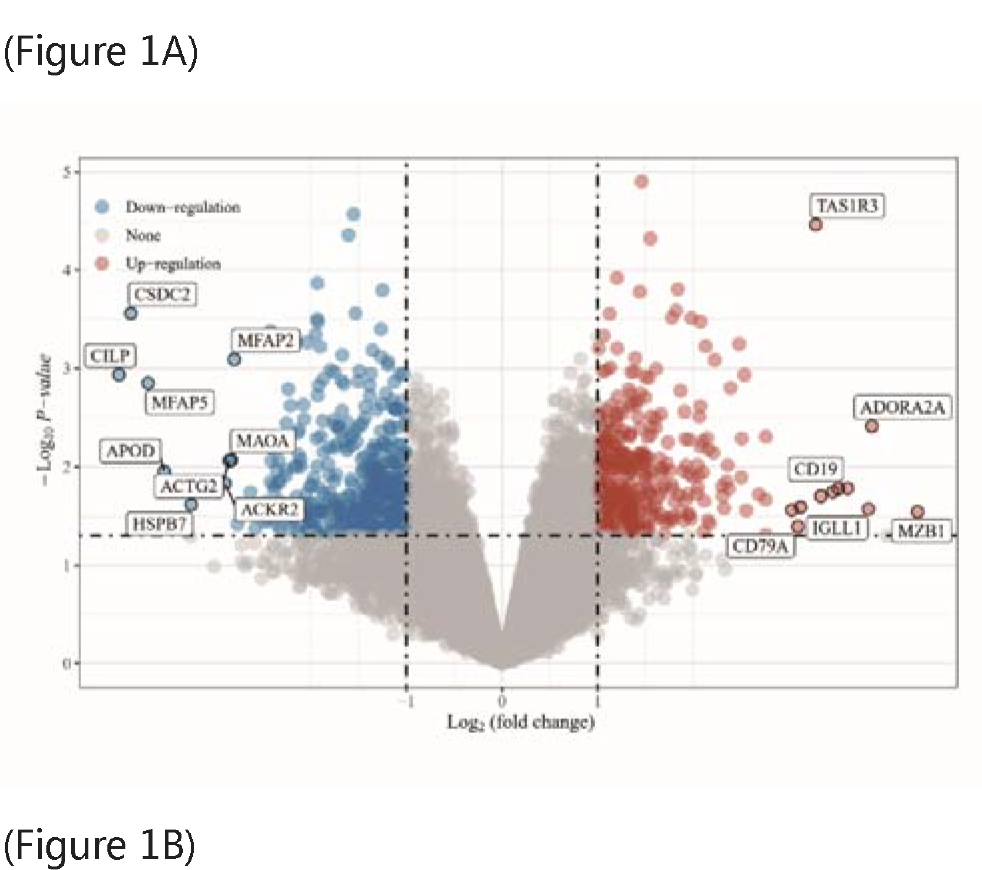
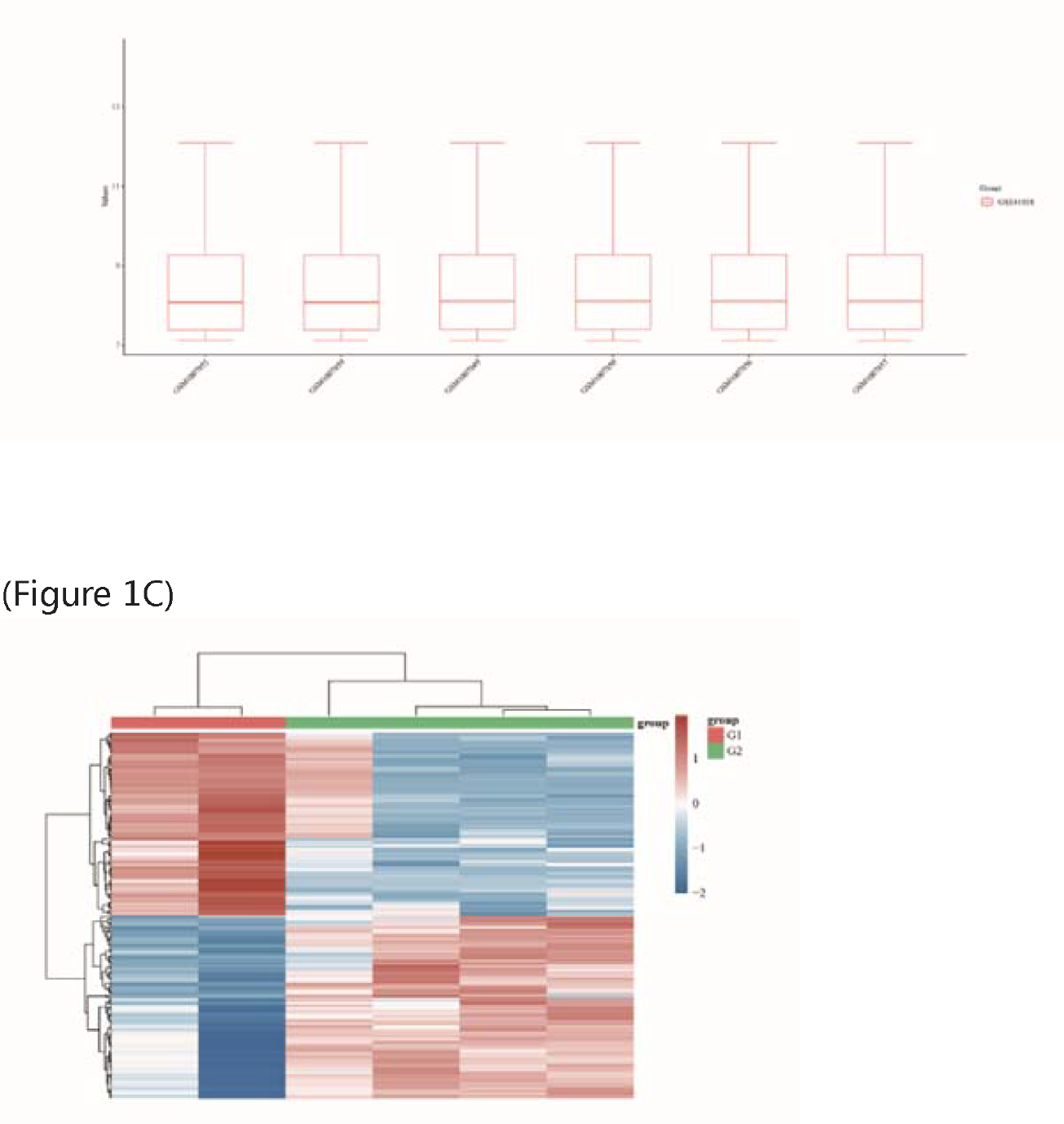
Gene expression of ankylosing spondylitis. (A) Volcanic plots of gene expression of AS. Red represents up-regulated DEGs, blue represents down-regulated DEGs, and grey represents genes that are not differentially expressed. (B) Box plots indicate the cross-comparability of the selected samples (C) Heatmap of the DEGs.

### Ferroptosis-Related Genes and Venn Analysis

From the iron death database (http://www.zhounan.org/ferrdb/), 259 Ferroptosis-related and 20 overlapping genes were obtained by taking intersections with differential genes, as demonstrated in the Venn diagram (Figure 2A). The iron death-related modular genes were: ASNS, GCH1, BACH1, RIPK1, WIPI1, SLC1A4, PLIN2, DUSP1, FADS2, NQO1, SCD, DUOX1, ZEB1, AKR1C2, CDO1, DPP4, AKR1C3, NOX4, GPT2, and CAV1). The sample distribution of the 20 genes is displayed in Figure 2B, and the correlation analysis is illustrated in Figure 2C.

**Figure 2.**
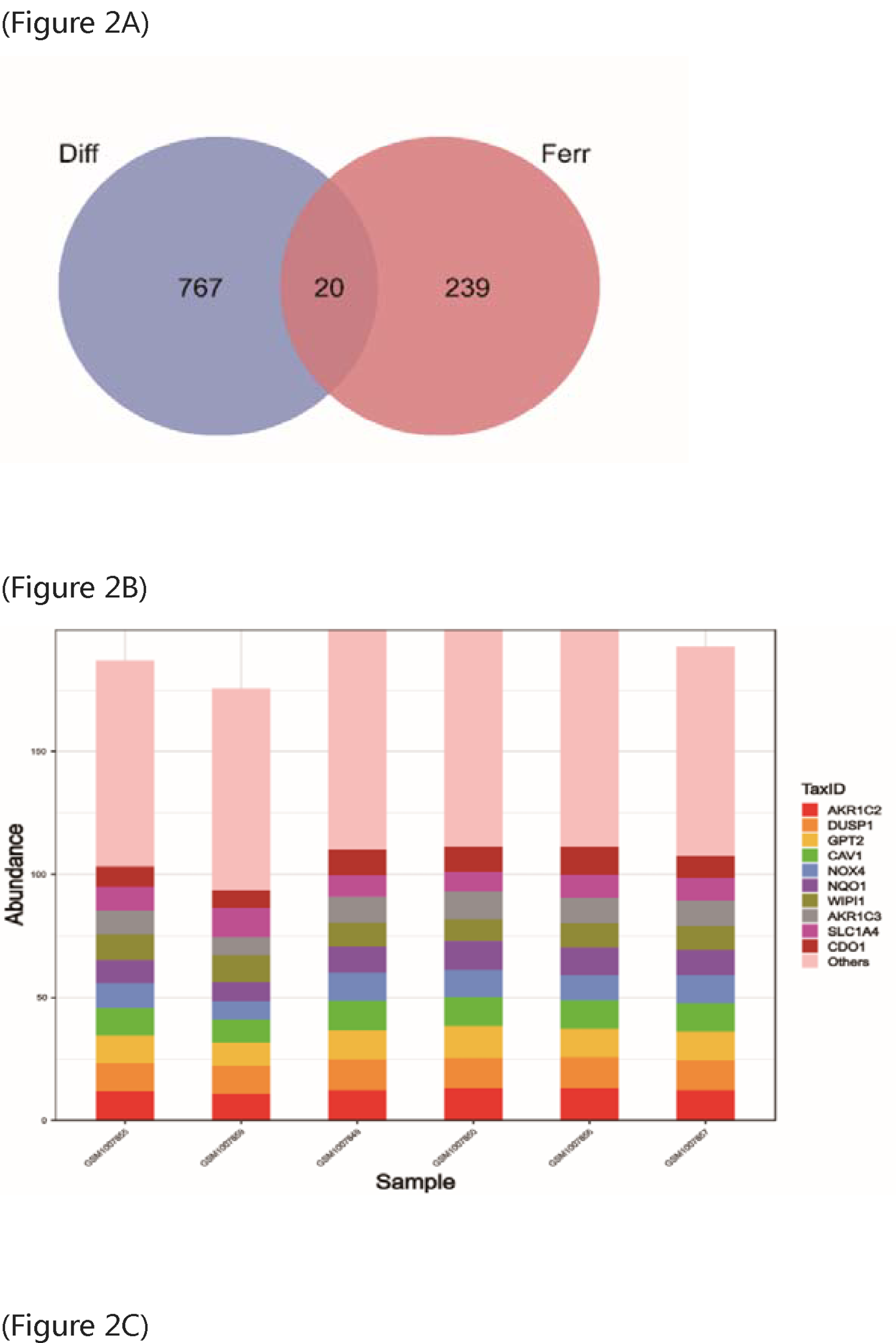
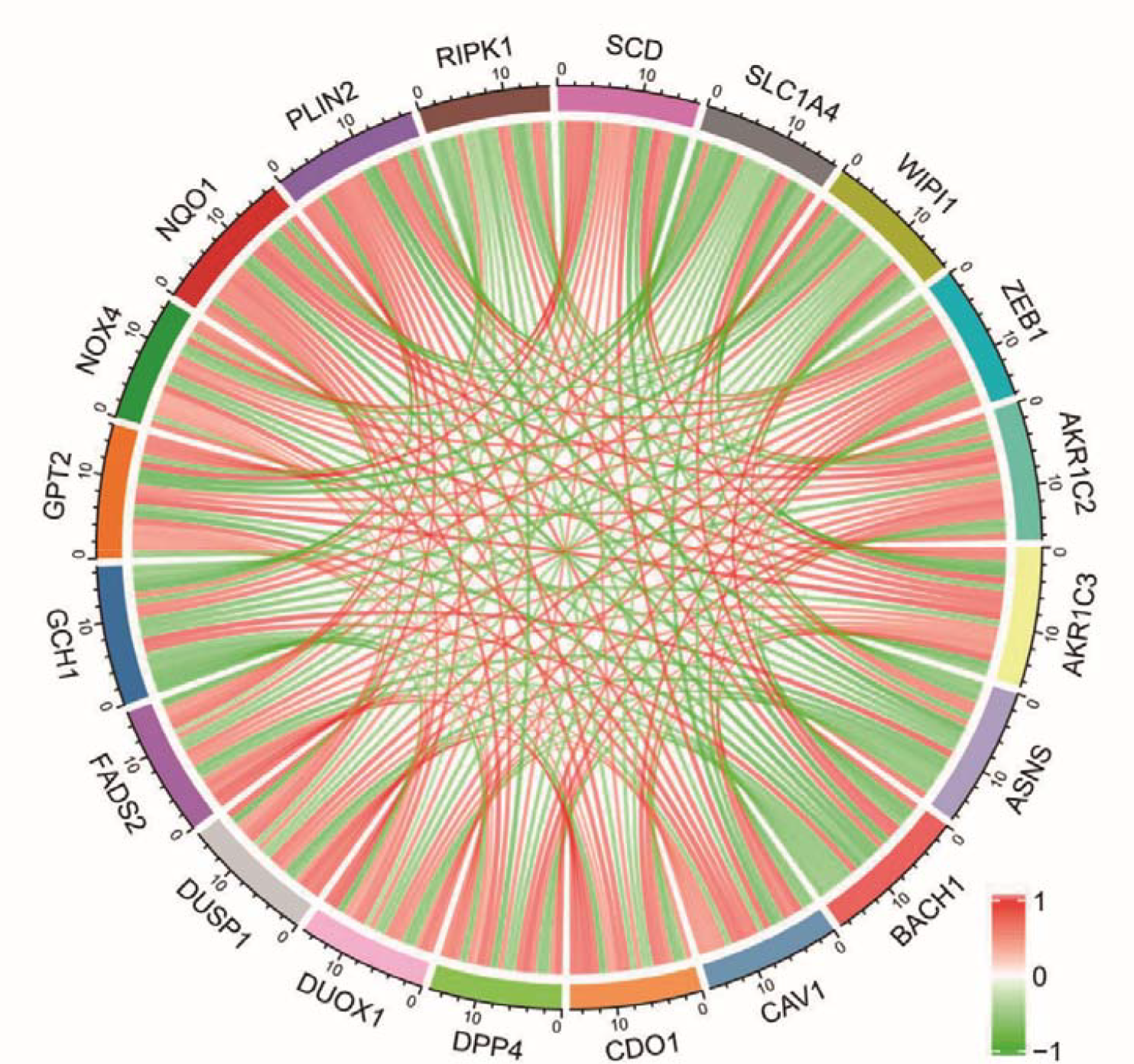
Analysis of Differentially expressed gene (DEG) in AS. (A) Venn diagram of ferroptosis-related DEGs, (B) Sample distribution of 20 genes, (C) Correlation analysis of 20 genes.

### Gene Ontology (GO) Enrichment and Kyoto Encyclopedia of Genes and Genome (KEGG) Pathway Analysis

The gene ontology enrichment analysis of ferroptosis-related differential genes consisted of three parts: biological process, cellular component, and molecular function. For biological processes, the ferroptosis-related DEGs were primarily enriched in the regulation of reactive oxygen species metabolic process, carboxylic acid biosynthetic process, and organic acid biosynthetic process (Figure 3A). The significantly enriched cellular component included the membrane raft, the apical part of the cell, and the apical plasma membrane (Figure 3B). The most enriched molecular functions were oxidoreductase activity, NAD(P)H activity, oxidoreductase activity, paired donors activity, incorporation or reduction of molecular oxygen, and steroid binding (Figure 3C). The KEGG enrichment results indicated that the ferroptosis-related DEGs were predominantly enriched in Chemical carcinogenesis-reactive oxygen species, Alcoholic liver disease, and PPAR signaling pathway (Figure 3D).

**Figure 3.**
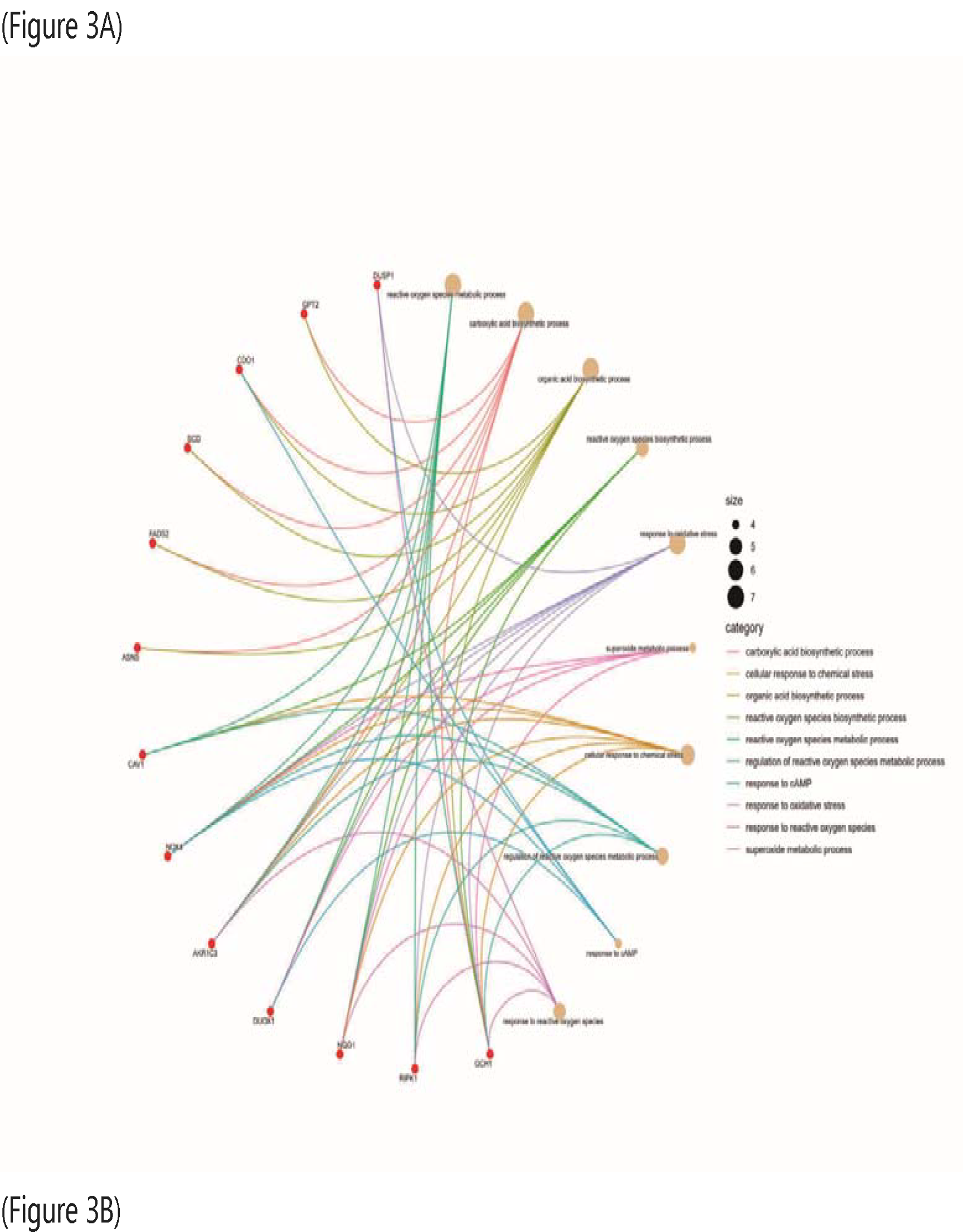
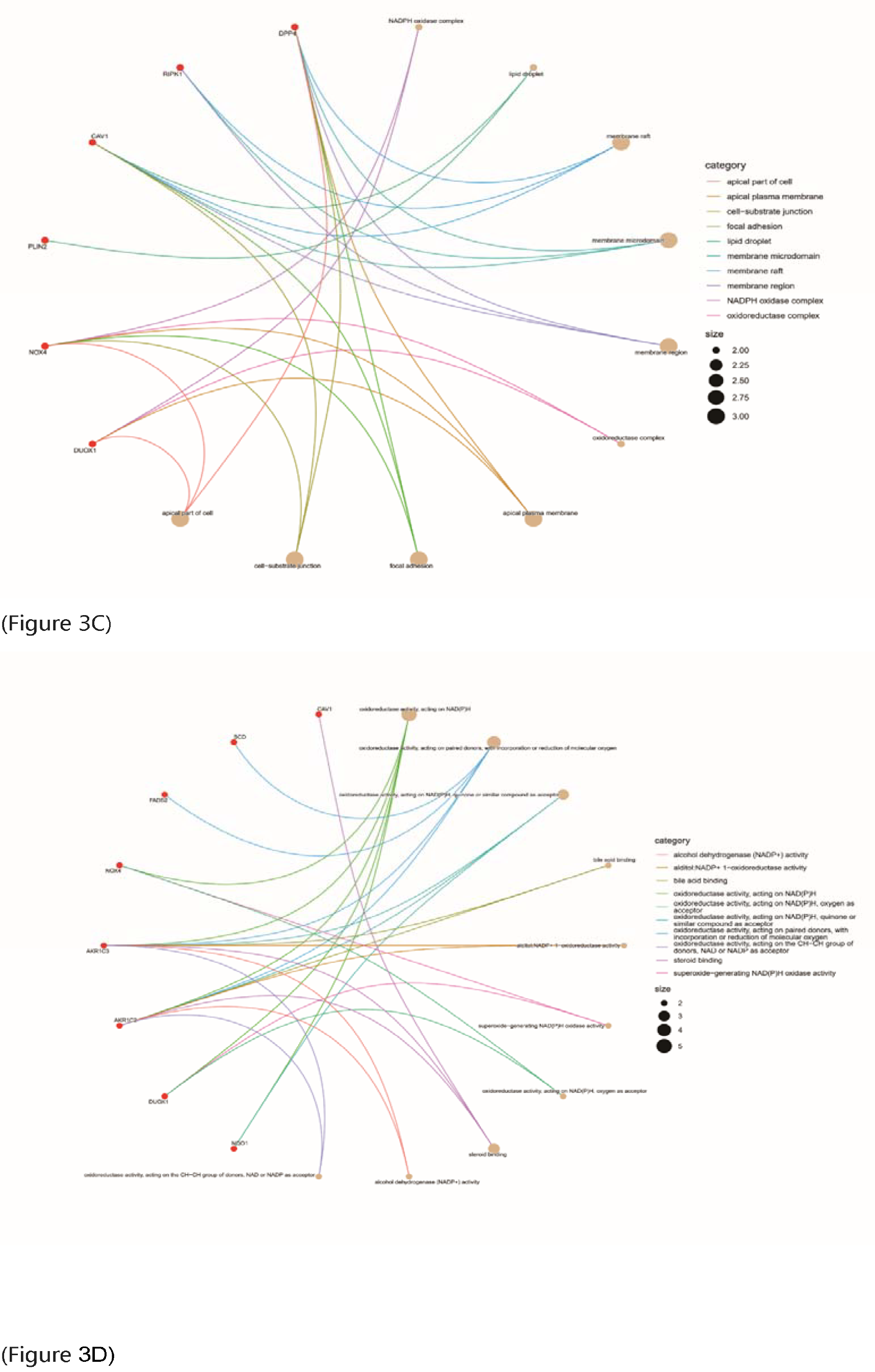
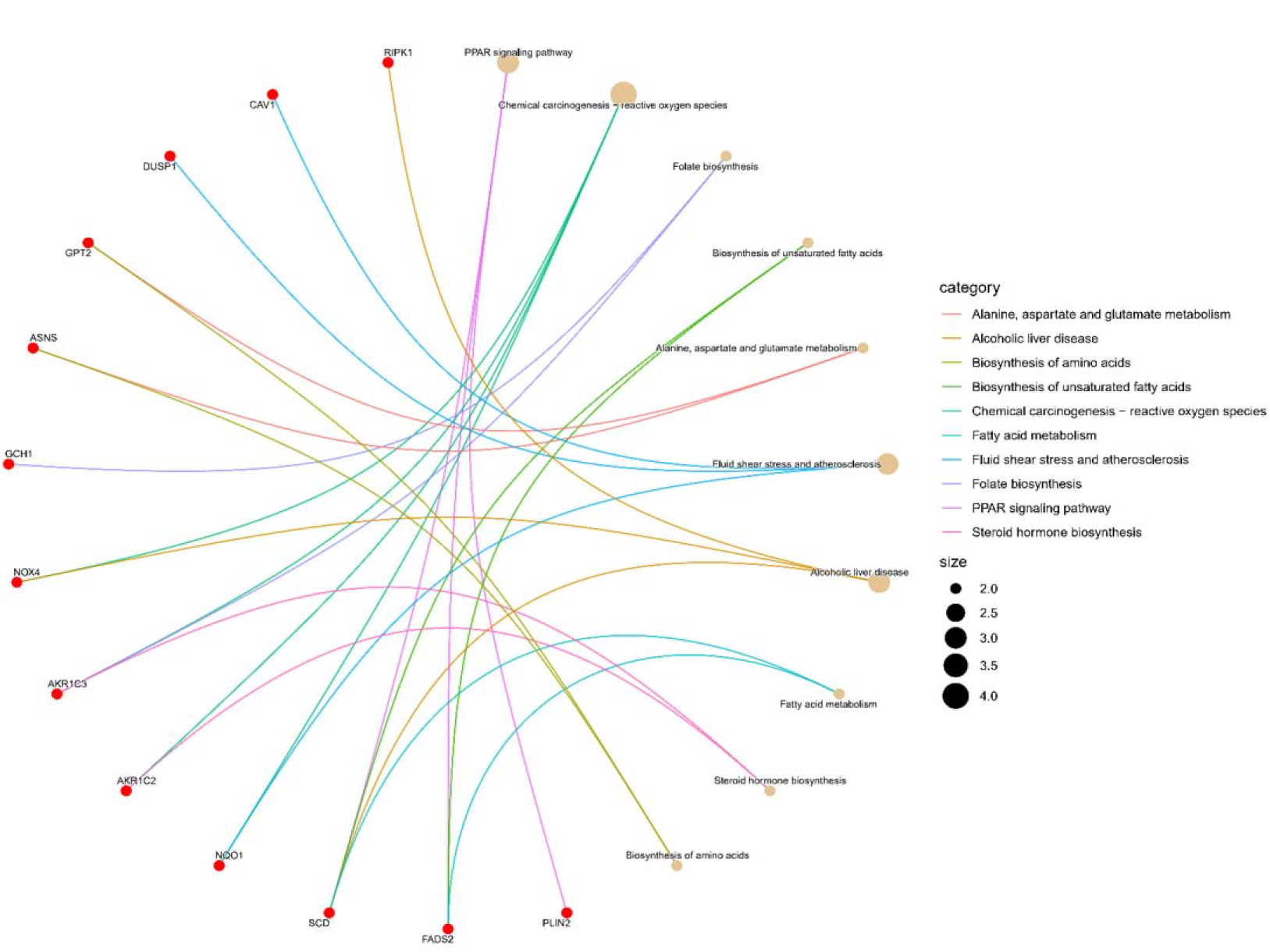
Enrichment analysis of the differentially expressed genes between the two clusters, using Gene Ontology (GO) and Kyoto Encyclopedia of Genes and Genomes (KEGG). (ABC) The top 10 enriched GO terms in cellular components, molecular functions, and biological processes. (D) The top 10 enriched KEGG pathways.

### Protein-Protein Interaction Network Construction and Hub Genes Analysis

To further investigate the role of iron death-associated DEGs, we performed PPI analysis on 20 genes from STRING for protein-protein interactions to determine the interactions between differentially expressed iron death-associated genes. Based on these results, we constructed an ankylosing spondylitis network with 20 nodes and 14 edges (Figure 4A). In addition, we imported the PPI network into Cytoscape analysis (Figure 4B) using the MCC algorithm to calculate HUB genes (Figure 4C). The top 10 hub genes are listed in Table 1.

**Table1.** Top 10 hub genes.

| Rank | Name | Score |
| --- | --- | --- |
| 1 | CAV1 | 4 |
| 2 | NOX4 | 3 |
| 2 | NQO1 | 3 |
| 4 | DUOX1 | 2 |
| 4 | AKR1C3 | 2 |
| 4 | ASNS | 2 |
| 4 | AKR1C2 | 2 |
| 4 | FADS2 | 2 |
| 4 | PLIN2 | 2 |
| 4 | SCD | 2 |

**Figure 4.**
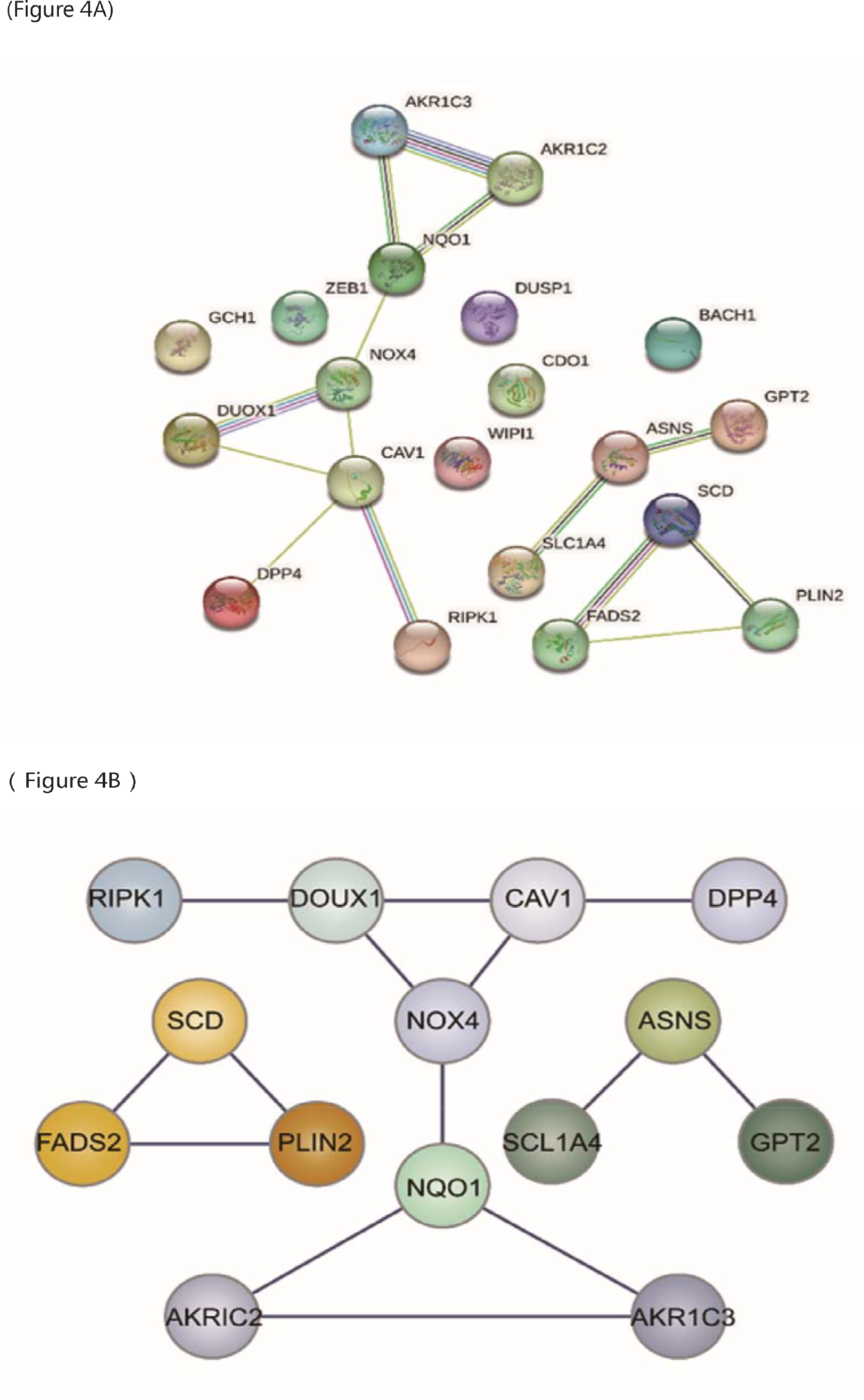
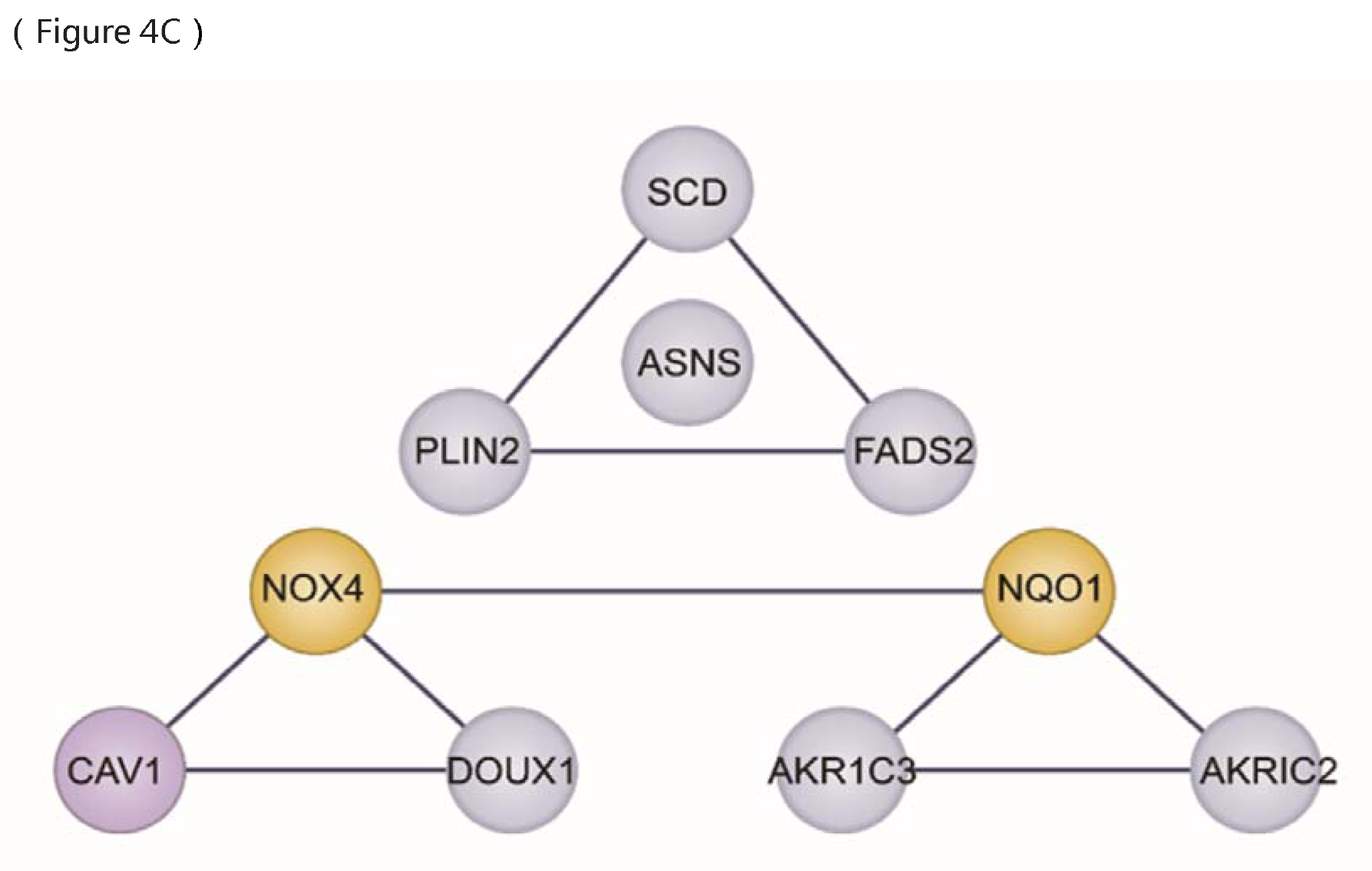
(A) The protein-protein interaction network constructed using STRING database representing the degree of gene interaction, (B) Importing PPI networks into Cytoscape for analysis(4C)Top 10 hub genes.

### Machine learning model building and evaluation

The neural network and C5.0 decision tree model used the presence or absence of AS as the output variable and the remaining 20 genes as the input variables (Figure 5A). The data node is the data source for this study, the Type is the node description for the variables, and the Partition node in this study is to randomly partition the samples into training and validation sample sets in a 3:1 ratio. At the Partition node, the random number of seeds was set to 1234567 so that random sample partition results could be repeated. The accuracy of the neural network model for 20 genes (training set: prediction set = 3:1) is 100% (Figure 5B).

**Figure 5.**
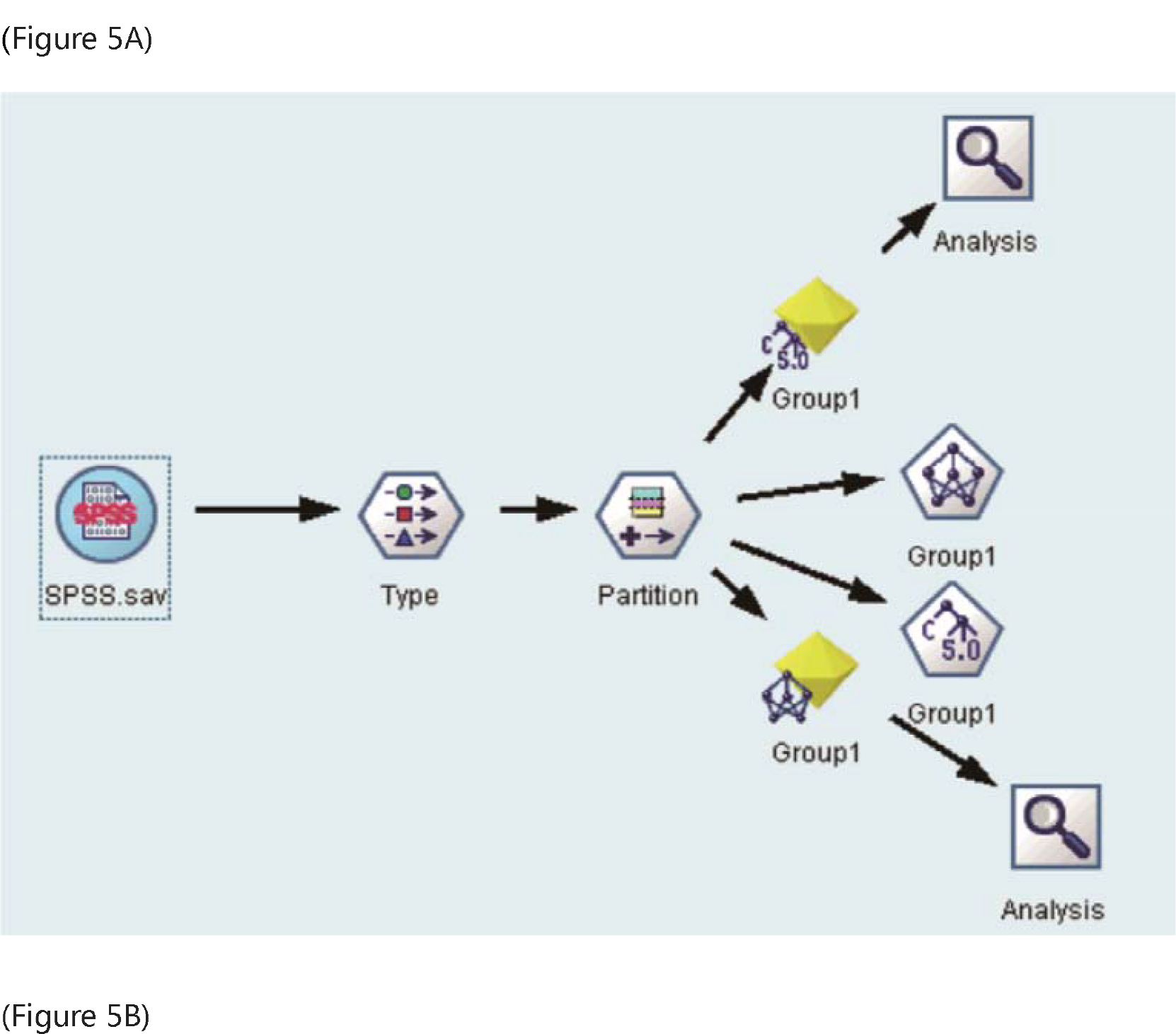
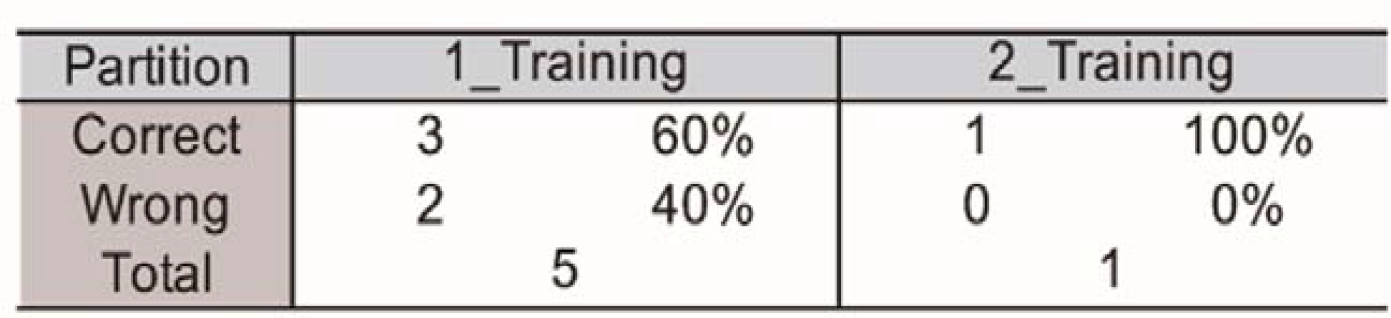
(A) Machine learning model building, (B) Neural Network model results for 20 genes.

### Screening for Potential Pharmacological Targets

The screening results for potential pharmacological targets downloaded from cMAP were ranked and filtered based on the linkage score of the drugs. The top 10 drugs suggested for AS were NCF2, TBX2, PP-30, ADAM15, SLC2A6, HSP90AB1, entinostat, entinostat, COBL, and ochratoxin-a (Table 2). These pharmaceuticals may have a therapeutic effect on AS. According to the results of screening the target genes of AS and 10 predicted drugs based on the CTD database, a total of 7421 target genes were identified for AS. A total of 2735 target genes for the predicted drug entinostat and 4 target genes for KU-0063794 were identified. The intersection was then utilized to generate a Cytoscape-based network pharmacology map of the disease, target genes, and drugs (Figure 6).

**Table 2.** Top 10 prediction results from cMap for AS.

| Score | ID | Name | Description |
| --- | --- | --- | --- |
| -99.19 | CGS001-4688 | NCF2 | Tetratricopeptide (TTC) repeat domain-containing |
| -98.47 | CGS001-6909 | TBX2 | T-boxes |
| -98.38 | BRD-K30677119 | PP-30 | RAF inhibitor |
| -98.3 | CGS001-8751 | ADAM15 | ADAM metallopeptidase domain containing |
| -98.28 | CGS001-11182 | SLC2A6 | Solute carriers |
| -97.63 | CGS001-3326 | HSP90AB1 | Heat shock proteins / HSPC |
| -97.36 | BRD-K77908580 | entinostat | HDAC inhibitor |
| -97.28 | BRD-K67566344 | entinostat | MTOR inhibitor |
| -96.61 | CGS001-23242 | COBL | - |
| -96.18 | BRD-K39944607 | ochratoxin-a | Phenylalanyl tRNA synthetase inhibitor |

**Figure 6.**
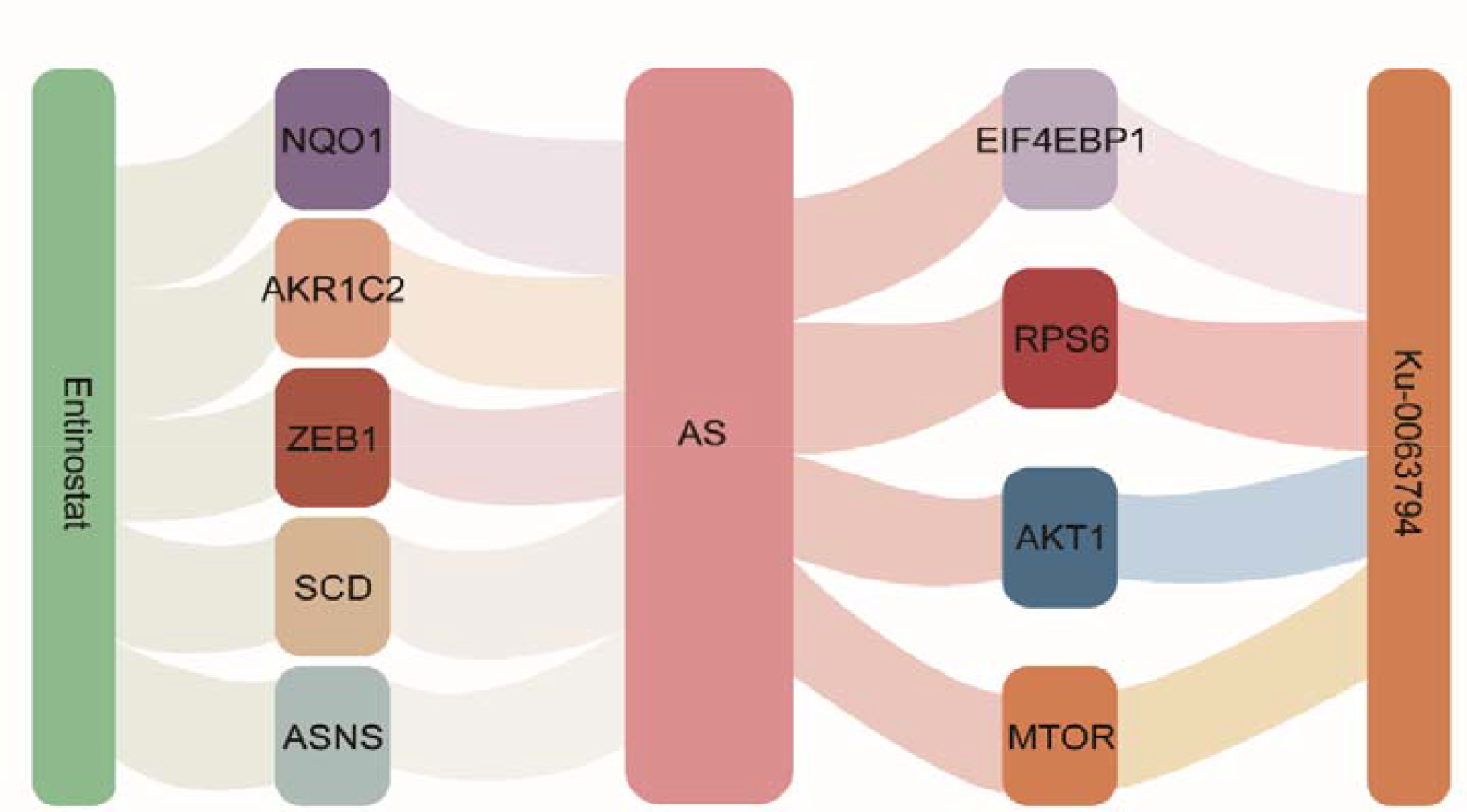
Pharmacological network map of predictive drugs entinostat and KU-0063794 with AS.

## DISCUSSION

The pathogenesis of AS is poorly understood, and the genes and pathways associated with Ferroptosis in AS have yet to be identified. Ankylosing spondylitis is an immune-mediated inflammatory arthritis member of a larger group of spondyloarthropathies (SpA) that also includes reactive arthritis, enteropathic arthritis, and psoriatic arthritis. AS predominantly affects the medial skeleton, particularly the sacroiliac and spinal joints, resulting in severe chronic pain and disability ^8^. The previous reliance on X-rays for the diagnosis of medial spondyloarthritis, magnetic resonance imaging (MRI) and HLA-B27 ^15^ allow us to diagnose this disease before it progresses to its tonic form and to initiate treatment earlier for better long-term outcomes. In the present study, we first analyzed the gene expression profiles of AS patients to identify differentially expressed genes associated with iron death. The identified differentially expressed genes in AS suggests that iron death-related genes are involved in the pathogenesis of AS. To understand the role of these Ferroptosis related genes in AS, we performed GO enrichment analysis and KEGG Pathway Analysis, which revealed the principal biological processes and signaling pathways involved in the content (Figure 3).

KEGG pathway analysis revealed that iron-death-associated differentially expressed genes in AS are primarily involved in the PPAR signaling pathway. PPARs are nuclear receptors and transcription factors consisting of PPARα, PPARβ/δ and PPARγ. These transcription factors regulate genes involved in lipid oxidation ^16^. It has been reported that in treating AS, Huangqin Qingre Chubi Capsules (HQC) decreased all disease activity index scores, increased mRNA levels of PPARγ, and increased protein levels of PPARγ ^17^. In an experiment investigating the effect of serum on the classical Wnt/β-linked protein pathway in ankylosing spondylitis patients, increased PPARD expression was observed in the serum of AS patients ^18^. These findings support the use of targeted therapies for AS.

Ten genes central to iron death in AS identified by Cytoscape were: CAV1, NOX4, NQO1, DUOX1, AKR1C3, ASNS, AKR1C2, FADS2, PLIN2, and SCD (Table 2). The Cav1 channel family, also known as L-type Ca2+ channels (LTCC), is extremely sensitive to organic Ca2+ channel blockers and is expressed in numerous electrically excited tissues ^19^. NOX4 is a constitutively active enzyme that primarily produces hydrogen peroxide and exhibits properties completely distinct from those of other NOX family isoforms, expressing in lungs, ovary, pancreas, and other organs ^20, 21^. However, the specific mechanism of NOX4 in the development of AS is unclear, and this study provides new insights into the NOX4 treatment of AS. NQO1 is associated with diabetes and metabolic syndrome prevention ^22^. Diaz-Ruiz et al. found that NQO1-deficient mice exhibit reduced fasting insulin levels and improved glucose metabolism ^23^. Dual oxidase 1 (DUOX1) is a member of the Nicotinamide Adenine Dinucleotide Phosphate (NADPH) oxidases protein family. Human DUOX1 is highly expressed in the lungs, pancreas, placenta, prostate, testis, and salivary glands ^24^. Human aldo-keto reductase family 1 member C3 (AKR1C3) regulates the occupancy of hormone receptors and cell proliferation as a prostaglandin F (PGF) synthase ^25^. AKR1C2 is essential for androgen conversion in adipose tissue ^26^. Asparagine synthetase ASNS is present in most mammalian organs, and mutations in the ASNS gene are clinically associated with asparagine synthetase deficiency (ASD), which can result in developmental delay and intellectual disability ^27^ and the development of cancer ^28^. Fatty acid desaturase 2 (Fads2) is the essential enzyme for the biosynthesis of long-chain polyunsaturated fatty acid (LC-PUFA) ^29^. Studies have shown that PLIN2 is implicated in the intracellular mobilization and storage of neutral lipids ^30^. Plin2 appears to be the most distinctive protein associated with fatty liver disease ^31^. The results of screening the target genes of AS and 10 predictive drugs using the CTD database identified a total of 74enes for AS. A total of 2735 target genes for the predicted drug entinostat and 4 target genes for KU-0063794 were identified.

There are several limitations to this study. First, when selecting datasets for differential expression analysis, the number of datasets and associated AD patients that could be selected remained constrained, and it was dis. It was discovered that some datasets contained few or no differentially expressed genes. In addition, if DEGs overlap with iron death-related module genes, the number of available genes is reduced and cannot be used for further analysis. Most of the data collected to date are limited to detecting changes in gene expression and require validation through additional functional experiments.

## CONCLUSION

The results of this study suggest that iron death genes may play a significant role in the pathogenesis and molecular regulation of ankylosing spondylitis. The key genes and pathways involved were identified using bioinformatic analysis and the drugs for the treatment of the disease treating. These findings contribute to our understanding of AS and its treatment.

## Data availability statement

Publicly available datasets were analyzed in this study. This data can be found here: https://www.ncbi.nlm.nih.gov/geo/query/acc.cgi?acc=gse41038

## Authors’ contributions

Zhen-Gang Liu and Fan Yang performed data analyses and wrote the manuscript. Bing Chen and Peng-Fu Li analyzed and interpreted data. Bo-Yin Zhang reviewed the first draft and gave comments.

## Declarations

### Consent for publication

This article has not been published or accepted for publication.

### Consent to participate

All authors have read and approved the manuscript

## Funding

This study was funded by the Provincial Natural Fund of Science and Technology Department of Jilin Province(20200201436JC), Jilin University Bethune Program (2023B22)

Jilin University Research Program(2022YX0256)

## REFERENCES

1. Fuchs Y, Steller H. Programmed cell death in animal development and disease. Cell. 2011;147(4): 742–758. 10.1016/j.cell.2011.10.033.

2. Qiu Y, Cao Y, Cao W, Jia Y, Lu N. The Application of Ferroptosis in Diseases. Pharmacological research. 2020;159: 104919. 10.1016/j.phrs.2020.104919.

3. Dixon SJ, Lemberg KM, Lamprecht MR, et al. Ferroptosis: an iron-dependent form of nonapoptotic cell death. Cell. 2012;149(5): 1060–1072. 10.1016/j.cell.2012.03.042.

4. Weiland A, Wang Y, Wu W, et al. Ferroptosis and Its Role in Diverse Brain Diseases. Molecular neurobiology. 2019;56(7): 4880–4893. 10.1007/s12035-018-1403-3.

5. Son E, Lee D, Woo CW, Kim YH. The optimal model of reperfusion injury in vitro using H9c2 transformed cardiac myoblasts. The Korean journal of physiology & pharmacology : official journal of the Korean Physiological Society and the Korean Society of Pharmacology. 2020;24(2): 173–183. 10.4196/kjpp.2020.24.2.173.

6. Feng H, Stockwell BR. Unsolved mysteries: How does lipid peroxidation cause ferroptosis? PLoS biology. 2018;16(5): e2006203. 10.1371/journal.pbio.2006203.

7. Ebrahimiadib N, Berijani S, Ghahari M, Pahlaviani FG. Ankylosing Spondylitis. Journal of ophthalmic & vision research. 2021;16(3): 462–469. 10.18502/jovr.v16i3.9440.

8. Voruganti A, Bowness P. New developments in our understanding of ankylosing spondylitis pathogenesis. Immunology. 2020;161(2): 94–102. 10.1111/imm.13242.

9. Rosenbaum J, Chandran V. Management of comorbidities in ankylosing spondylitis. The American journal of the medical sciences. 2012;343(5): 364–366. 10.1097/MAJ.0b013e3182514059.

10. Yang H, Chen Y, Xu W, et al. Epigenetics of ankylosing spondylitis: Recent developments. International journal of rheumatic diseases. 2021;24(4): 487–493. 10.1111/1756-185x.14080.

11. Chen B, Li J, He C, et al. Role of HLA-B27 in the pathogenesis of ankylosing spondylitis (Review). Molecular medicine reports. 2017;15(4): 1943–1951. 10.3892/mmr.2017.6248.

12. Thomas GP, Duan R, Pettit AR, et al. Expression profiling in spondyloarthropathy synovial biopsies highlights changes in expression of inflammatory genes in conjunction with tissue remodelling genes. BMC musculoskeletal disorders. 2013;14: 354. 10.1186/1471-2474-14-354.

13. Zhou N, Bao J. FerrDb: a manually curated resource for regulators and markers of ferroptosis and ferroptosis-disease associations. Database : the journal of biological databases and curation. 2020;2020. 10.1093/database/baaa021.

14. Chin CH, Chen SH, Wu HH, Ho CW, Ko MT, Lin CY. cytoHubba: identifying hub objects and sub-networks from complex interactome. BMC systems biology. 2014;8 Suppl 4(Suppl 4): S11. 10.1186/1752-0509-8-s4-s11.

15. Zhu W, He X, Cheng K, et al. Ankylosing spondylitis: etiology, pathogenesis, and treatments. Bone research. 2019;7: 22. 10.1038/s41413-019-0057-8.

16. Contreras AV, Torres N, Tovar AR. PPAR-α as a key nutritional and environmental sensor for metabolic adaptation. Advances in nutrition (Bethesda, Md). 2013;4(4): 439–452. 10.3945/an.113.003798.

17. Huang D, Liu J, Zong RK, Wan L. [Huangqin Qingre Chubi Capsules in improving oxidative stress of patients with ankylosing spondylitis via activating PPAR? mediated AMPK/FOXO3a pathway]. Zhongguo Zhong yao za zhi = Zhongguo zhongyao zazhi = China journal of Chinese materia medica. 2020;45(2): 451–456. 10.19540/j.cnki.cjcmm.20190619.501.

18. Hu Z, Lin D, Qi J, et al. Serum from patients with ankylosing spondylitis can increase PPARD, fra-1, MMP7, OPG and RANKL expression in MG63 cells. Clinics (Sao Paulo, Brazil). 2015;70(11): 738–742. 10.6061/clinics/2015(11)04.

19. Striessnig J, Bolz HJ, Koschak A. Channelopathies in Cav1.1, Cav1.3, and Cav1.4 voltage-gated L-type Ca2+ channels. Pflugers Archiv : European journal of physiology. 2010;460(2): 361–374. 10.1007/s00424-010-0800-x.

20. Hiraga R, Kato M, Miyagawa S, Kamata T. Nox4-derived ROS signaling contributes to TGF-β-induced epithelial-mesenchymal transition in pancreatic cancer cells. Anticancer research. 2013;33(10): 4431–4438.

21. Carnesecchi S, Deffert C, Donati Y, et al. A key role for NOX4 in epithelial cell death during development of lung fibrosis. Antioxidants & redox signaling. 2011;15(3): 607–619. 10.1089/ars.2010.3829.

22. Martínez-Hernández A, Córdova EJ, Rosillo-Salazar O, et al. Association of HMOX1 and NQO1 Polymorphisms with Metabolic Syndrome Components. PloS one. 2015;10(5): e0123313. 10.1371/journal.pone.0123313.

23. Diaz-Ruiz A, Di Francesco A, Carboneau BA, et al. Benefits of Caloric Restriction in Longevity and Chemical-Induced Tumorigenesis Are Transmitted Independent of NQO1. The journals of gerontology Series A, Biological sciences and medical sciences. 2019;74(2): 155–162. 10.1093/gerona/gly112.

24. Edens WA, Sharling L, Cheng G, et al. Tyrosine cross-linking of extracellular matrix is catalyzed by Duox, a multidomain oxidase/peroxidase with homology to the phagocyte oxidase subunit gp91phox. The Journal of cell biology. 2001;154(4): 879–891. 10.1083/jcb.200103132.

25. Liu Y, He S, Chen Y, et al. Overview of AKR1C3: Inhibitor Achievements and Disease Insights. Journal of medicinal chemistry. 2020;63(20): 11305–11329. 10.1021/acs.jmedchem.9b02138.

26. Zhang Y, Dufort I, Rheault P, Luu-The V. Characterization of a human 20alpha-hydroxysteroid dehydrogenase. Journal of molecular endocrinology. 2000;25(2): 221–228. 10.1677/jme.0.0250221.

27. Lomelino CL, Andring JT, McKenna R, Kilberg MS. Asparagine synthetase: Function, structure, and role in disease. The Journal of biological chemistry. 2017;292(49): 19952–19958. 10.1074/jbc.R117.819060.

28. Dufour E, Gay F, Aguera K, et al. Pancreatic tumor sensitivity to plasma L-asparagine starvation. Pancreas. 2012;41(6): 940–948. 10.1097/MPA.0b013e318247d903.

29. Bláhová Z, Harvey TN, Pšenička M, Mráz J. Assessment of Fatty Acid Desaturase (Fads2) Structure-Function Properties in Fish in the Context of Environmental Adaptations and as a Target for Genetic Engineering. Biomolecules. 2020;10(2). 10.3390/biom10020206.

30. Brasaemle DL, Barber T, Wolins NE, Serrero G, Blanchette-Mackie EJ, Londos C. Adipose differentiation-related protein is an ubiquitously expressed lipid storage droplet-associated protein. Journal of lipid research. 1997;38(11): 2249–2263.

31. Okumura T. Role of lipid droplet proteins in liver steatosis. Journal of physiology and biochemistry. 2011;67(4): 629–636. 10.1007/s13105-011-0110-6.

